# Benchmarking of Hi-C tools for scaffolding de novo genome assemblies

**DOI:** 10.1101/2023.05.16.540917

**Authors:** Lia Obinu, Urmi Trivedi, Andrea Porceddu

**Affiliations:** Department of Agricultural Sciences, University of Sassari, Viale Italia 39A,07100, Sassari, Sardinia, Italy; Edinburgh Genomics, Ashworth Laboratories, Charlotte Auerbach Road, The King’s Buildings, The University of Edinburgh, EH9 3FL, Edinburgh, Scotland, United Kingdom

## Abstract

The implementation of Hi-C reads in the *de novo* genome assembly allows to order large regions of the genome in scaffolds, obtaining chromosome-level assemblies. Several bioinformatics tools have been developed for genome scaffolding with Hi-C, and all have pros and cons which need to be carefully evaluated before adoption.

We developed assemblyQC, a bash pipeline that combines QUAST, BUSCO, Merqury and, optionally, Liftoff, plus a gene positioning validation script to evaluate and benchmark the performance of three scaffolders, 3d-dna, SALSA2, and YaHS, on two de novo assembly of Arabidopsis thaliana obtained from the same raw PacBio HiFi and ONT data.

In our analysis, YaHS proved to be the best-performing bioinformatic tool for scaffolding of *de novo* genome assembly.

## Introduction

Third-generation sequencing technologies (TGS), e.g., Pacific Biosciences (PacBio) and Oxford Nanopore Technologies (ONT), produce long reads which allow to obtain high-quality genome sequences. Compared to next-generation sequencing technologies, TGS give better resolution and contiguity, allowing to partially resolve the assembly of repeats and duplications, which are crucial to resolve repetitive rich genomic regions such as the centromeres and telomeres, especially in plant genomes [1].

Although long read assembly on itself can yield long contigs, this can still be far from assembling to a chromosome-scale without additional scaffolding efforts. Strategically, larger genome projects like the Vertebrate Genome Project (VGP - https://vertebrategenomesproject.org/), and The Darwin Tree of Life (DToL - https://www.darwintreeoflife.org/) combine multiple data types like optical mapping [2] and Hi-C [3] to construct reliably chromosome-scale assemblies [4]. Originally developed to study the three-dimensional organisation of the genome [5] Hi-C reads are now largely employed for scaffolding *de novo* genome assemblies [6]. Hi-C technology is a method which combines proximity-based ligation with massively parallel sequencing allowing the unbiased identification of chromatin interactions across an entire genome. This allows to scaffold together large regions of the genome, obtaining chromosome-level assemblies [7].

A previous study conducted a benchmarking of Hi-C-based scaffolders partitioned reference genomes and *de novo* assemblies [8]. The scaffolders’ performance evaluation was based on the mapping of assemblies against the known reference genome using Mummer 4 [9], and on a set of accuracy metrics calculated using the Python package Edison (https://github.com/Noble-Lab/edison).

In this study, we benchmarked three Hi-C scaffolders (3d-dna [10], SALSA2 [11], and YaHS [12]) starting from two *de novo* assemblies of *A. thaliana*, the model species for plant genomics [13], obtained from the same PacBio Hifi and ONT raw data. 3d-dna was chosen for being the most applied Hi-C scaffolder still under active development [8]. SALSA2 is the new version of SALSA [14], while YaHS is the most recently released scaffolder. Indeed, the last two scaffolders were never benchmarked with other scaffolders before.

To perform the benchmarking, we developed assemblyQC, a bash pipeline that combines QUAST [15], BUSCO [16], and Merqury [17]. The pipeline also optionally uses Liftoff [18] to annotate the *de novo* assembly, and a Python script subsequently analyses the positioning of genes on the target assembly compared to the reference.

## Materials and methods

### *Arabidopsis thaliana* genome and raw data

The first *A. thaliana* genomic sequence was obtained in 2000 from the Columbia genotype, using the method of the minimum tiling path of BACs sequenced with Sanger’s technology [19]. The genome sequence length is approximately 135 Mb in length, organised in 5 chromosomes (2n = 10).

In this study, we used the publicly available raw data from the BioProject PRJCA005809 [20] which was later used to build two *de novo* assemblies and evaluate the performance of the three scaffolders thereafter.

Table 1 summarises the characteristics of the raw reads used. Illumina reads were not used for the assembly process, but they were used with Merqury [17] for scaffolding evaluation purposes.

**Table 1:** Overview of the characteristics of the raw reads from the BioProject PRJCA005809 [20] used in the assembly process.

| Reads | Mean read length | Mean read quality | Number of reads | Total bases | Coverage |
| --- | --- | --- | --- | --- | --- |
| ONT | 18,541.30 | 11.1 | 3,064,191.00 | 56,814,196,989.00 | 420.84507 |
| PacBio HiFi | 15,094.40 | 31.8 | 1,517,433.00 | 22,904,700,074.00 | 169.66475 |
| Hi-C | 150 | >30 | 140,957,500 | 21,143,625,000 | 156.6194444 |
| Illumina | 150 | >30 | 91,309,542 | 13,696,431,300 | 101.4550467 |

### De novo assembly

ONT reads were trimmed with NanoFilt [21] using the parameter -l 500, which only retained the reads with a minimum length of 500 bp.

The first *de novo* assembly was obtained by assembling ONT reads using Flye [22] in --nano-raw mode with default parameters. To subsequently perform the polishing, the PacBio HiFi reads were mapped to the draft assembly using minimap2 [23] in map-hifi mode, and then the polishing was undertaken with Racon [24]. Haplotigs and overlaps in the assembly were removed using purge_dups [25]. Contaminants were removed using BlobToolKit [26] applying a filter for GC content equal to 0.4 since Tiara [27] identified contigs with higher GC content as “unknown”. Hereafter, we will refer to this assembly as “Flye”.

The second *de novo* assembly was obtained by assembling HiFi and ONT reads together using hifiasm [28] with default parameters. Haplotigs and overlaps in the assembly were removed using purge_dups [25]. Contaminants were removed using BlobToolKit [26] applying a filter for GC content equal to 0.5 since Tiara [27] identified contigs with higher gc content as “unknown” or “bacteria”. Furthermore, a filter for HiFi coverage equal to 5180 was applied as contigs above this level of coverage were identified as organelles by Tiara. Hereafter, we will refer to this assembly as “Hifiasm”.

### Hi-C scaffolding

All the Hi-C scaffolders combine Hi-C linkage information with draft genome assemblies to resolve contig orientations.

The 3d-dna algorithm as a first step identifies assembly errors where a scaffold’s long-range contact pattern changes unexpectedly. Thereafter the resulting sequences are anchored, ordered and oriented thanks to an algorithm based on the contact frequency between pairs of sequences as an indicator of their proximity in the genome. Finally, contigs and scaffolds that correspond to overlapping regions of the genome are merged together [10].

SALSA2 builds a scaffold graph scoring edges according to a “best buddy” scheme. The scaffolds are then constructed according to a greedy weighted maximum matching. Afterwards, SALSA2 performs an iterative step of mis-join detection and correction, which stops naturally when accurate Hi-C links are exhausted [11].

YaHS is the newest scaffolder that has been developed and, compared to the previous ones, it proposes a new method for building the contact matrix. The software proceeds building a contact matrix, constructing and pruning a scaffold graph, and giving out the scaffolds [12].

Hi-C reads were trimmed before performing the scaffolding using fastp with the following parameters: `-p, --detect_adapter_for_pe, --cut_front, --cut_tail, --cut_window_size 4, --cut_mean_quality` 20.

### 3d-dna

Hi-C reads were filtered and aligned to the draft assembly, using Juicer [29] with parameters `-p assembly, -s none, -S early`. The fasta file of the contig-level assembly was given as input for both the Juicer flags --g, normally used to input a reference assembly, and `--z`, to avoid reference biases. Prior to scaffolding, the contig-level assembly was wrapped using the script `wrap-fasta-sequence.awk`. Finally, the run-asm-pipeline.sh script from 3d-dna [10] was run with the wrapped. `fasta` file and the list of Hi-C contacts in. `txt` format outputted by Juicer. The script used were those provided for single CPU run.

### SALSA2

Hi-C reads were filtered and aligned to the contig-level assembly following the Arima Genomics mapping pipeline (https://github.com/ArimaGenomics/mapping_pipeline). The steps 1A and 1B of the pipeline were modified adding the flag `-M` to the bwa mem commands. The resulting bam file was subsequently sorted by read name using `samtools sort` and converted in .bed file using `bedtools`. Finally, the script `run_pipeline.py` from SALSA2 [11] was run with parameters `-e GATC` and `-m` yes to perform the scaffolding.

### YaHS

The Hi-C reads were filtered and aligned to the contig-level assembly following the same methods described for SALSA2. The output `.bam` file of the Arima Genomics mapping pipeline was sorted and converted to .bed format as described above. YaHS was finally run with parameters `-e GATC` [12].

### AssemblyQC pipeline

We developed assemblyQC, a Bash pipeline designed to perform the quality control of the assemblies. AssemblyQC combines QUAST, BUSCO, and Merqury, and optionally runs Liftoff along with a Python script that produces metrics about gene positioning in the assembly compared to a given reference genome.

QUAST [15] is a tool which computes relevant quality metrics useful to evaluate *de novo* assemblies and compare them against reference sequences. Using the flag `-w` assemblyQC runs QUAST-LG, the QUAST extension used for evaluating large genomes. In our case, regular QUAST was run.

BUSCO [16] evaluates genome assembly quality in terms of gene completeness. To run BUSCO using assemblyQC it is necessary to previously download the database of the lineage of interest. In our case this was `brassicales_odb10`.

Merqury [17] allows to perform reference-free assembly evaluation of accuracy and completeness, comparing the k-mers found in the *de novo* assembly with those present in high accuracy raw reads. In order to run Merqury, it’s necessary to previously generate a k-mer database using Meryl. In our case, Illumina short reads were used to build the Meryl database. AssemblyQC supports only the case in which one assembly with no hap-mers (i.e. haplotype specific k-mers) is provided to run Merqury.

Liftoff [18] is a standalone tool that accurately maps annotation from a reference genome to a target assembly. It requires reference `.fasta` and `.gff/`.`gtf` files and the. `fasta` file of the target assembly; the output is a. `gff/.gtf` file for the target assembly. Assembly QC accepts as input for Liftoff only files in. `gff` format, and optionally filters out organelles’ annotations before running this program.

AssemblyQC runs Liftoff with the flag `--exclude-partial`, used to exclude partial/low sequence identity mappings (coverage >= 0.5 and sequence identity >= 0.5) from the output `.gff`.

`liftoff_combine`.py is a python script we developed to output metrics about gene collinearity between the target assembly and the reference genome. We define gene collinearity as the correspondence between the gene positioning in the target assembly compared to the reference genome. The script takes the output `.gff` from Liftoff and the reference `.gff` file as input. It only analyses gene collinearity, therefore exons and transcripts are excluded from the. `gff` files. Moreover, the script allows to set a threshold to evaluate the divergence in intergenic length between pairs of adjacent genes between the target and the reference assemblies: if the divergence is lower than the threshold, the compared genomic annotation are considered coherent (default: 500 bp).

`liftoff_combine.py` outputs a `.txt` file per reference chromosome that gives information regarding: i) which contigs or scaffolds in the target assembly correspond to the reference chromosome; ii) in which orientation contigs or scaffolds in the target assembly are compared to the reference; iii)if and in which measure the distance between adjacent genes in the target assembly match the distance between corresponding genes in the reference assembly.

Secondary outputs of this script are a merged assembly-reference `.gff` for whole genome, merged `.gff` files per chromosome, and records of the genes showing divergent gene distance between the reference and the assembly per chromosome.

For this study, we ran assemblyQC including the Liftoff step. We removed organelles annotations from the reference `.gff` file and adopted a divergence threshold of 500 bp for `liftoff_combine.py`. The input k-mer length was determined with the Merqury command `best_k.sh`.

### Evaluation criteria

The evaluation of the two scaffolded assemblies was based on several approaches: 1) the QUAST metrics of genome fraction, number of contigs, N50, N90, L50, and L90 and the cumulative length plot; 2) the metrics provided by BUSCO (Complete BUSCOs, Complete and single-copy BUSCOs, Complete and duplicated BUSCOs, Fragmented BUSCOs, and Missing BUSCOs) comparing the assemblies with the database for Brassicales (`brassicales_odb10`); 3) Merqury consensus quality (QV), k-mer completeness and copy number spectrum plots; 4) gene collinearity metrics provided by the Liftoff plus `liftoff_combine.py` method; 5) Hi-C contact map generated using JuiceBox Assembly Tools (JBAT) [30].

### Cluster characteristics

The programmes in benchmarking were executed on a linux machine containing two 2.1 GHz, 18-core Intel Xeon E5-2695 (Broadwell) series processors where each of the cores in these processors support 2 hyperthreads enabled by default with 256 GB of memory.

## Results and discussion

### QUAST based benchmarking

Table 2 shows the QUAST metrics of the scaffolded assemblies compared to the contig-level assemblies Flye and Hifiasm. All the scaffolders were able to align more than 99% of bases to the reference genome of *A. thaliana* (TAIR10.1).

**Table 2:** Comparison of the assemblies according to QUAST metrics.

| Assembly | Genome fraction (%) | Contigs | N50 | N90 | L50 | L90 |
| --- | --- | --- | --- | --- | --- | --- |
| flye | 99.097 | 15 | 14,864,979 | 9,471,025 | 4 | 8 |
| flye_3d-dna | 99.068 | 30 | 19,600,500 | 12,160,500 | 3 | 6 |
| flye_salsa | 99.105 | 14 | 15,405,308 | 9,471,025 | 4 | 8 |
| flye_yahs | 99.109 | 7 | 24,256,539 | 19,109,354 | 3 | 5 |
| hifiasm | 99.302 | 5 | 26,162,003 | 22,263,862 | 3 | 5 |
| hifiasm_3d-dna | 99.304 | 342 | 3,413,500 | 175,000 | 8 | 139 |
| hifiasm_salsa | 99.302 | 8 | 16,698,585 | 13,446,829 | 4 | 7 |
| hifiasm_yahs | 99.287 | 4 | 32,656,027 | 22,674,124 | 2 | 4 |

SALSA2 and YaHS were able to reduce the number of contigs of the draft assembly, while 3d-dna increased it. The best result was achieved by YaHS for both the assemblies. YaHS reduced the number of contigs from 15 to 7 in the case of Flye, and from 5 to 4 in the case of Hifiasm, probably introducing a false joint between two contigs as 5 chromosomes are expected for *A. thaliana*. Notably, the number of contigs increased dramatically for Hifiasm scaffolded with 3d-dna, rising up to 342 contigs from the initial 5 contigs.

All the scaffolders increased the N50 compared to the draft assemblies, except for Hifiasm scaffolded with 3d-dna, where the N50 decreased from the initial ∼26 Mb to ∼3 Mb. Only YaHS increased the N90 for both Flye (from ∼9 Mb to ∼19 Mb) and Hifiasm (from ∼22.3 Mb to ∼22.7 Mb). 3d-dna increased the N90 only for Flye (from ∼9 Mb to ∼12 Mb), while it strongly decreased the N90 in the case of Hifiasm (from ∼22 Mb to ∼2Mb). SALSA2 didn’t produce changes in N90 in the case of Flye, while it decreased it for Hifiasm (from ∼22 Mb to ∼13 Mb). YaHS produced both the highest N50 and N90 for both the assemblies.

Flye L50 was reduced from 4 to 3 by both 3d-dna and YaHS, while no L50 changes were produced by SALSA2. Hifiasm L50 was decreased from 3 to 2 by YaHS. YaHS also achieved the lowest L90 for both the assemblies.

The full QUAST report is shown in supplementary materials (Table S1).

Figure 1 shows the QUAST plot for cumulative length (Mbp). Flye assembly was improved by all the scaffolders, with YaHS giving the best results for all the parameters considered. In this case, 3d-dna slightly increased the fragmentation of the assembly increasing the number of contigs, but it improved all the other parameters. SALSA2 reduced the number of contigs and increased the N50, keeping the other parameters unchanged.

**Figure 1:**
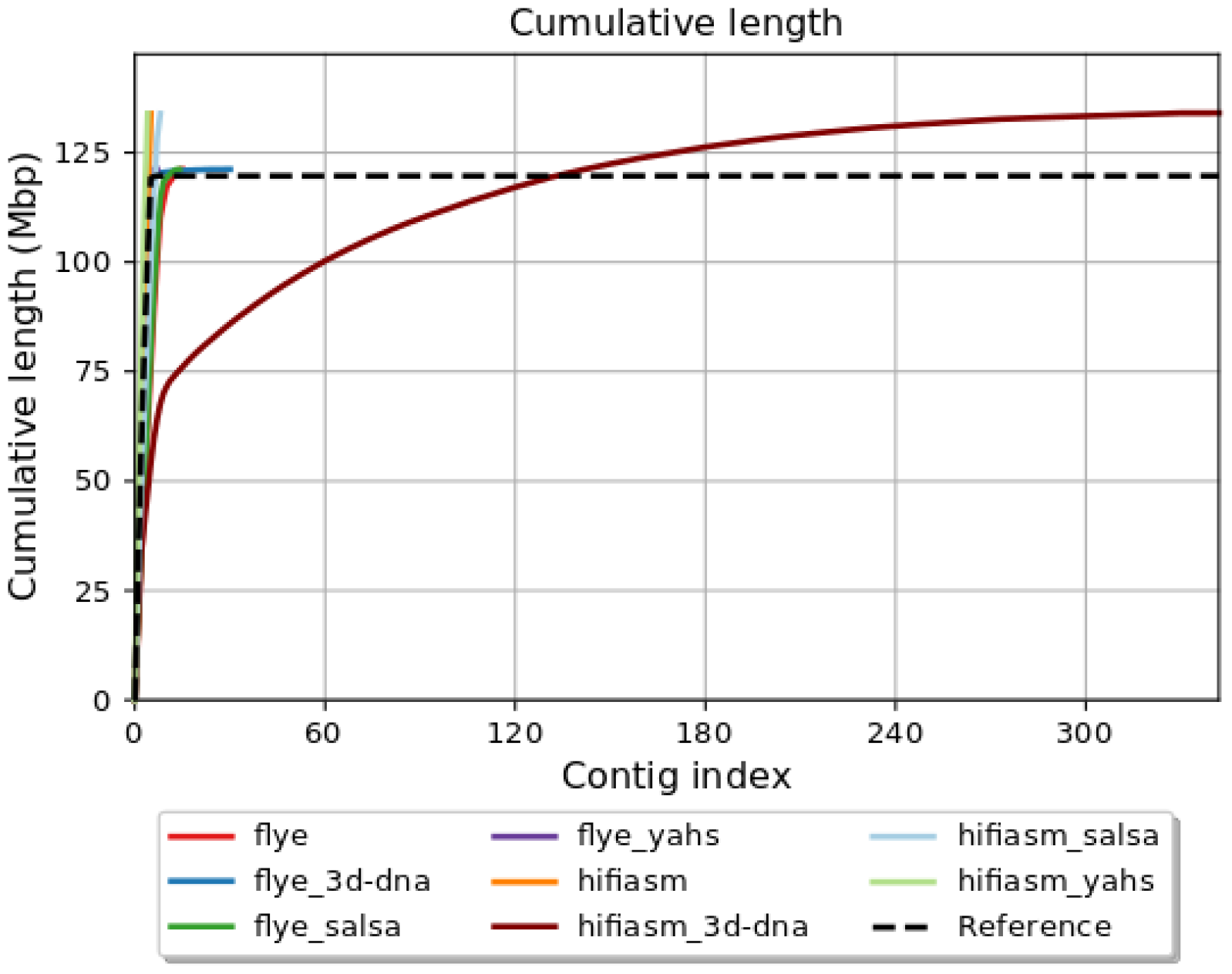
QUAST plot for cumulative length (Mbp).

In conclusion, Hifiasm assembly showed good metrics already at contig-level. YaHS was the only scaffolder that didn’t increase the assembly fragmentation, but it introduced a false joint, joining two contigs in the same scaffold. SALSA2 and 3d-dna increased the assembly fragmentation, with the latter representing the most extreme case.

### BUSCO based benchmarking

Figure 2 shows the comparison between Flye and Hifiasm and the relative scaffolded assemblies as regard the BUSCO metrics.

**Figure 2:**
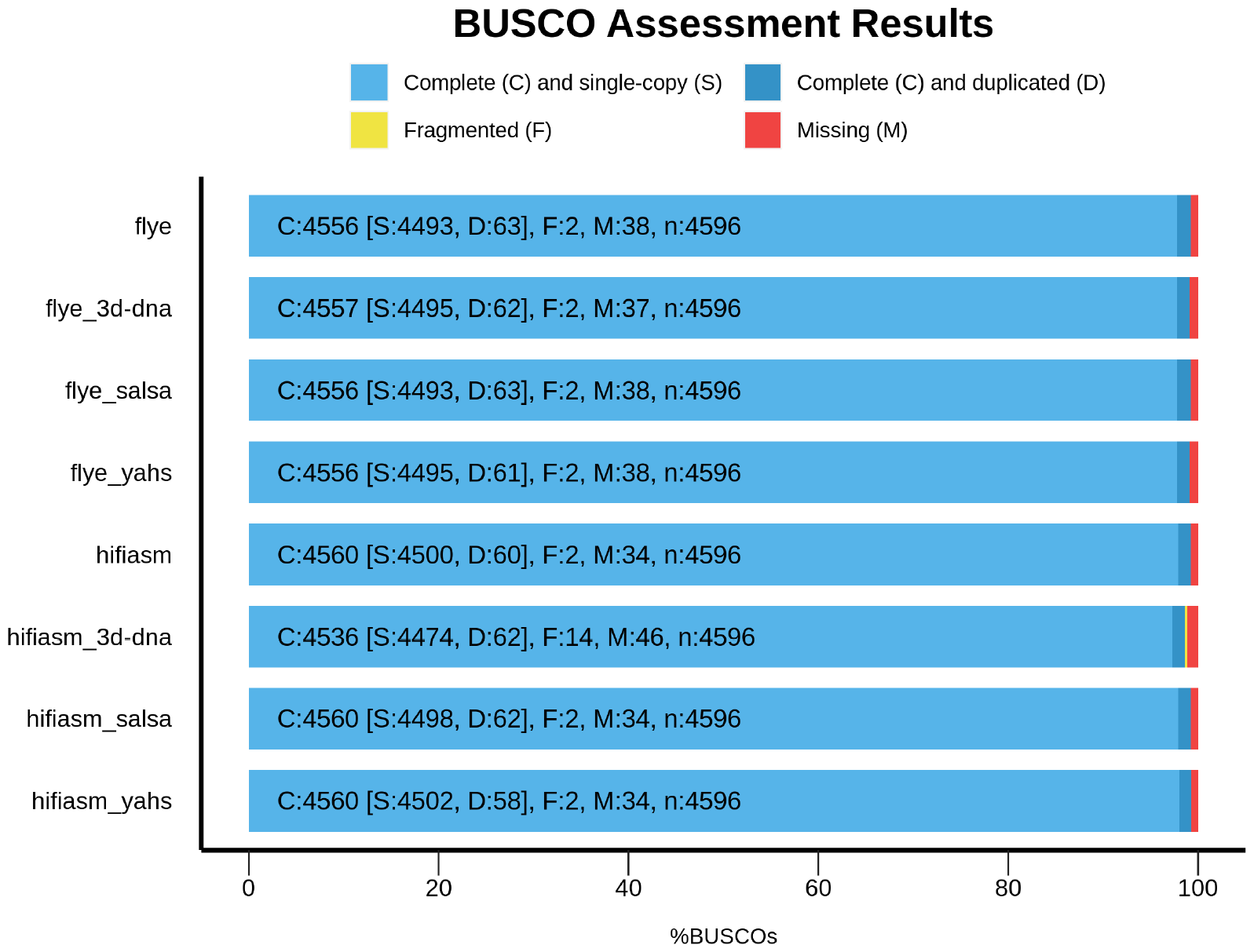
Comparison of the assemblies according to BUSCO metrics.

SALSA2 and YaHS kept the number of complete, missing and fragmented BUSCOs unchanged compared to the contig-level assemblies.

3d-dna behaved differently according to the assembly considered. In the case of Flye, it increased the number of complete BUSCOs, it decreased the number of missing BUSCOs and kept fragmented BUSCOs unaltered compared to the contig-level assembly, overall improving the BUSCO metrics. On the other hand, in the case of Hifiasm 3d-dna decreased the number of complete BUSCOs, and increased the number of missing and fragmented BUSCOs, overall decreasing the metrics quality compared to the contig-level assembly.

### Merqury based benchmarking

Table 3 shows the results obtained for each assembly with Merqury.

**Table 3:**
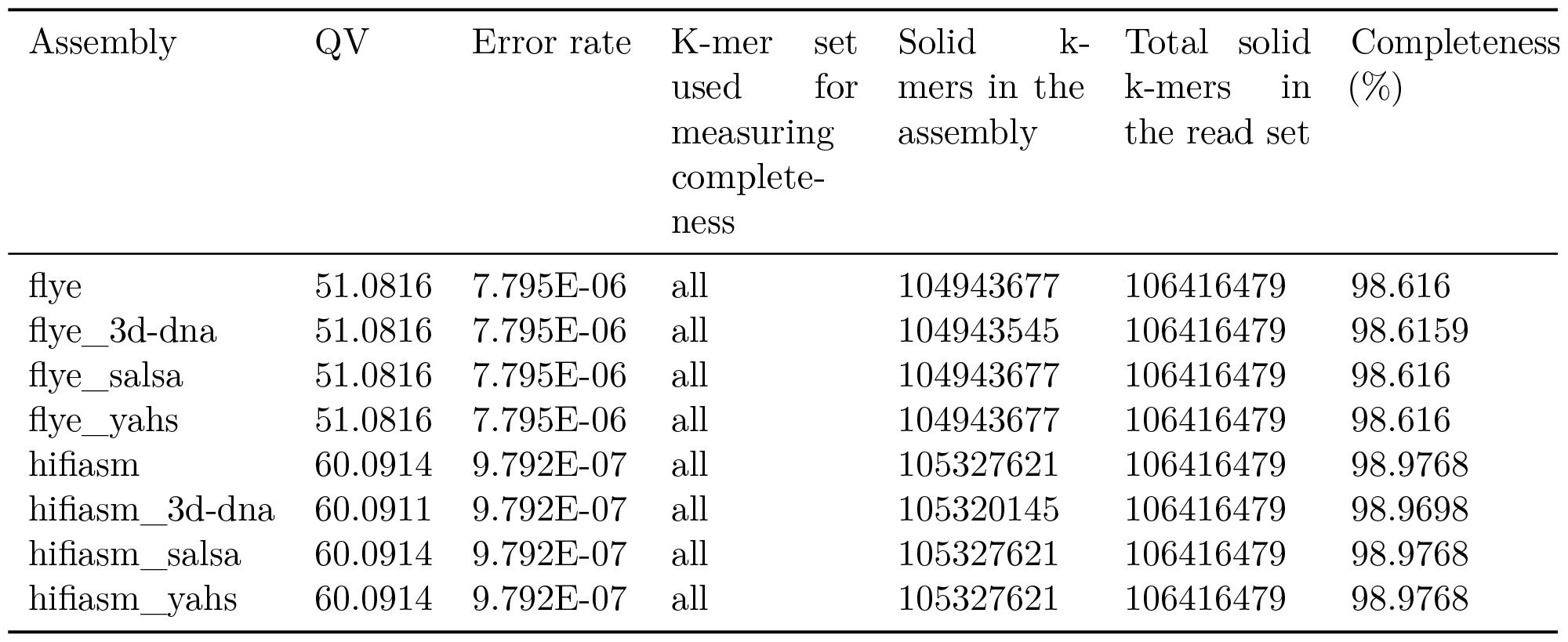
Comparison of the assemblies according to Merqury metrics.

The QV value in the scaffolded assemblies compared to the contig-level assemblies remained unaltered, except for Hifiasm scaffolded with 3d-dna, for which the QV value decreased from 60.0914 to 60.0911.

All the Flye assemblies showed a level of completeness above the 98.6%, and all the Hifiasm assemblies showed completeness >98.9%.

Copy number spectrum plots for each assembly are shown in supplementary materials (Figure S1). All the assemblies fit with the expected copy number spectrum for a haploid genome.

### Collinearity based benchmarking

Detailed results about the comparison of gene positioning in the assemblies and the reference genome TAIR10.1 are reported in supplementary materials (Table S2, S3, S4, S5, S6).

Table 4 displays the number of contigs or scaffolds in the target assemblies containing genes mapping to chromosome 1 to 5 in the reference genome TAIR10.1 according to Liftoff results.

**Table 4:** Number of contigs or scaffolds in the target assemblies containing genes annotated in chromosome 1 to 5 in the reference genome TAIR10.1 according to Liftoff results.

| Assembly | Chr 1 | Chr 2 | Chr 3 | Chr 4 | Chr 5 |
| --- | --- | --- | --- | --- | --- |
| Flye | 2 | 9 | 4 | 4 | 3 |
| Flye_3d-dna | 5 | 8 | 3 | 2 | 3 |
| Flye_salsa | 3 | 9 | 4 | 2 | 3 |
| Flye_yahs | 1 | 4 | 2 | 1 | 1 |
| Hifiasm | 1 | 1 | 3 | 1 | 1 |
| Hifiasm_3d-dna | 64 | 36 | 64 | 47 | 73 |
| Hifiasm_salsa | 2 | 2 | 3 | 1 | 2 |
| Hifiasm_yahs | 1 | 1 | 2 | 1 | 1 |

YaHS was the only scaffolder able to reduce the number of scaffolds corresponding to the TAIR10.1 chromosomes according to gene content in comparison to both the contig-level assemblies and for all the chromosomes.

Notably, as regards Hifiasm, which was initially formed by 5 contigs (i.e. the expected number of chromosomes), 3d-dna dramatically increased the number of scaffolds corresponding to the reference chromosomes according to gene content.

As regards the orientation of the scaffolds, we considered a contig or scaffold to have a clear orientation with respect to the reference genome when at least the 90% of the gene coordinates were in ascending (forward orientation) or descending order (reverse orientation).

As regards Flye, 3d-dna gave a clear orientation to all the chromosomes, except for the scaffolds corresponding to chromosome 2, where the proportion of genes in the same orientation as the reference and the proportion of genes in reverse complement orientation were 86.6% and 13.1 respectively. SALSA2 conserved the orientation of the scaffolds of the Flye contig-level assembly for chromosomes 1 and 3, which were in the same orientation and in reverse complement respectively compared to the reference genome. It wasn’t able to define a clear orientation for the scaffolds corresponding to chromosomes 2, 4 and 5 in TAIR10.1. YaHS was able to find a clear orientation for all the scaffolds.

In the case of Hifiasm, 3d-dna wasn’t able to define scaffolds orientation for any chromosome, except for the scaffolds corresponding to the chromosome 4 of TAIR10.1, which were in reverse complement orientation compared to the reference genome, as the contig-level Hifiasm assembly. SALSA2 found a clear orientation for the scaffolds corresponding to chromosome 1, 3, and 5, while their orientation is not clear for chromosomes 2 and 4. YaHS was able to find a clear orientation for all the scaffolds.

Overall, YaHS was the only scaffolder that for both the assemblies decreased the number of scaffolds corresponding to the reference chromosomes and found a clear orientation for all the scaffolds. The collinearity results confirmed that YaHS introduced a false joint in Hifiasm assembly merging the scaffolds that correspond to chromosomes 3 and 4 in TAIR10.1.

Hifiasm assembly deserves particular attention. This assembly contained only 5 contigs mainly corresponding to the chromosomes for of TAIR10.1 before scaffolding. The Hifiasm contigs ptg000001l_1 and ptg000002l_1 contained mainly genes corresponding to chromosome 2 and chromosome 4 of TAIR10.1, respectively. However, ptg000001l_1 and ptg000002l_1 contained also 7 and 2 genes, respectively, mapping on TAIR 10.1 chromosome 3. YaHS partially resolved these misplacements: only the 7 genes that were in contig ptg000001l_1 were placed in the scaffold_4 corresponding to chromosome 2 of TAIR10.1, while the other two genes originally mapping in ptg000002l_1 were correctly placed in the scaffold_1 corresponding to chromosome 3. SALSA2 did not resolve neither these misplacements. Due to the high level of fragmentation introduced by 3d-dna it was impossible to evaluate how it dealt with these misplacements (Table 5). Further data are needed to ascertain whether the 7 genes of Hifiasm contig ptg000002l_1 represent a misplacement or a true translocation event.

**Table 5:** Correspondence between reference chromosome 3, contigs in Hifiasm and scaffolds in Hifiasm scaffolded with SALSA2 and YaHS for the 9 genes involved in an observed misplacement.

| Gene ID | Reference chromosome | Hifiasm contig | SALSA2 scaffold | YaHS scaffold |
| --- | --- | --- | --- | --- |
| AT3G00650 | CP002686.1 | ptg000001l_1 | scaffold_8 | scaffold_4 |
| AT3G41762 | CP002686.1 | ptg000001l_1 | scaffold_8 | scaffold_4 |
| AT3G41761 | CP002686.1 | ptg000001l_1 | scaffold_8 | scaffold_4 |
| AT3G06345 | CP002686.1 | ptg000001l_1 | scaffold_8 | scaffold_4 |
| AT3G41768 | CP002686.1 | ptg000001l_1 | scaffold_8 | scaffold_4 |
| AT3G06355 | CP002686.1 | ptg000001l_1 | scaffold_8 | scaffold_4 |
| AT3G41979 | CP002686.1 | ptg000001l_1 | scaffold_8 | scaffold_4 |
| AT3G00660 | CP002686.1 | ptg000002l_1 | scaffold_2 | scaffold_1 |
| AT3G06365 | CP002686.1 | ptg000002l_1 | scaffold_2 | scaffold_1 |

Detailed correspondence chromosomes-contigs and chromosome-scaffolds is reported in supplementary materials (Table S2, S3, S4, S5, S6).

### Hi-C contact maps

JuiceBox Assembly Tools (JBAT) [30] allows to visualise the Hi-C contact map and to manually curate the scaffolded assemblies(Figure 3 A, C, D, F). SALSA2 provides a method that only allows to visualise the contact map in JBAT, but doesn’t allow the highlighting of contigs and scaffolds and therefore the manual curation of the assembly is not possible(Figure 3 B, E).

**Figure 3:**
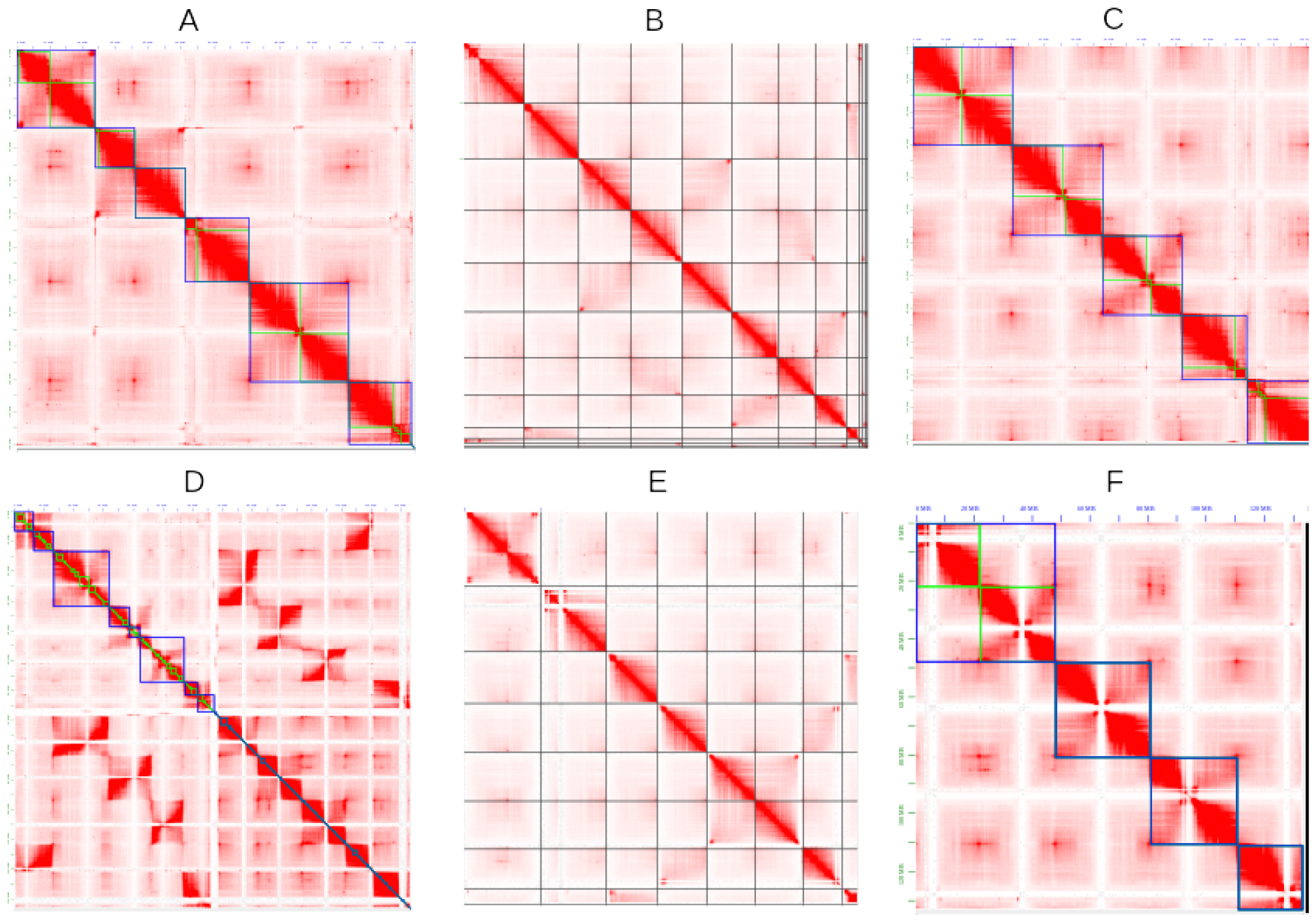
Contact maps obtained with JBAT. A, B and C show the contact map of the Flye assembly scaffolded with 3d-dna, SALSA2 and YaHS respectively. D, E and F show the contact map of the Hifiasm assembly scaffolded with 3d-dna, SALSA2 and YaHS respectively. In A, C, D and F green lines represent the contigs and blue lines the chromosome-length scaffolds.

The major scaffolds corresponding to the chromosomes were correctly identified in both Flye and Hifiasm only by YaHS (Figure 3C, 3F). In the last case, YaHS identified 4 major scaffolds, because two of the chromosome-length scaffolds were joined (Figure 3F). However, the error could be easily detected and corrected with JBAT, by manually splitting the two chromosome-length scaffolds.

The Hi-C contact map confirms that, for both the assemblies, 3d-dna and SALSA2 produced more fragmented assemblies compared to YaHS, with the most extreme case observed in Hifiasm scaffolded with 3d-dna.

## Conclusions

The two assemblies scaffolded with YaHS showed the best QUAST metrics, the most accurate gene positioning compared to the reference genome TAIR10.1, and the most accurate Hi-C contact map. YaHS was also the only tool to output chromosome length scaffolds with a clear orientation (either the same as the reference genome, or reverse complement) and to show them clearly in the Hi-C contact map.

SALSA2 and YaHS gave similar results as regard BUSCO metrics for both assemblies. 3d-dna improved BUSCO metrics for Flye assembly, but it gave poorer results for the Hifiasm assembly.

The assemblies scaffolded with SALSA2 showed the best QV according to Merqury results.

Overall, 3d-dna performed well with the Flye assembly, but it heavily fragmented the Hifiasm assembly, showing a possible incompatibility between the two tools.

YaHS was the easiest and quicker software to install and use, taking into account also the Hi-C pre-processing. In addition, the developers provide good documentation to implement the software in the analysis. 3d-dna also has good documentation, but we found that it is more difficult to install and run. As regard SALSA2, the main flaws are the lack of a detailed and rich documentation, and the method provided to visualise the contact map in JBAT, which doesn’t allow the manual curation of the assembly.

Of the three scaffolders that were benchmarked in this study, YaHS was the best performing for the majority of the parameters considered. Therefore, we conclude that it is the most appropriate scaffolding tool for *de novo* assemblies to date.

## Code availability

AssemblyQC is available at https://github.com/LiaOb21/assemblyQC.

## Acknowledgements

We would like to thank Dr. Alex Mackintosh of The University of Edinburgh for his useful advice and inputs on the topic.

