## Supplementary Materials for "Benchmarking of Hi-C tools for scaffolding de novo genome assemblies"

Table S1: Full QUAST report.

| Assembly | Flye | Flye_3d-dna | Flye_salsa | Flye_yahs | Hifiasm | Hifiasm_3d-dna | Hifiasm_salsa | Hifiasm_yahs |
|---|---|---|---|---|---|---|---|---|
| Contigs (≥ 0 bp) | 15 | 30 | 14 | 7 | 5 | 342 | 8 | 4 |
| Contigs (≥ 1000 bp) | 15 | 30 | 14 | 7 | 5 | 342 | 8 | 4 |
| Contigs (≥ 5000 bp) | 15 | 26 | 14 | 7 | 5 | 331 | 8 | 4 |
| Contigs (≥ 10000 bp) | 15 | 25 | 14 | 7 | 5 | 330 | 8 | 4 |
| Contigs (≥ 25000 bp) | 15 | 24 | 14 | 7 | 5 | 319 | 8 | 4 |
| Contigs (≥ 50000 bp) | 14 | 15 | 13 | 6 | 5 | 266 | 8 | 4 |
| Total length (≥ 0 bp) | 121168757 | 121178257 | 121169757 | 121169657 | 133901430 | 134049930 | 133902430 | 133901530 |
| Total length (≥ 1000 bp) | 121168757 | 121178257 | 121169757 | 121169657 | 133901430 | 134049930 | 133902430 | 133901530 |
| Total length (≥ 5000 bp) | 121168757 | 121168012 | 121169757 | 121169657 | 133901430 | 134034210 | 133902430 | 133901530 |
| Total length (≥ 10000 bp) | 121168757 | 121163012 | 121169757 | 121169657 | 133901430 | 134026210 | 133902430 | 133901530 |
| Total length (≥ 25000 bp) | 121168757 | 121152178 | 121169757 | 121169657 | 133901430 | 133826000 | 133902430 | 133901530 |
| Total length (≥ 50000 bp) | 121124559 | 120889192 | 121125559 | 121125459 | 133901430 | 132386186 | 133902430 | 133901530 |
| Contigs | 15 | 30 | 14 | 7 | 5 | 342 | 8 | 4 |
| Largest contig | 16316820 | 30170187 | 18141904 | 30360532 | 32656027 | 18736000 | 26162503 | 48425965 |
| Total length | 121168757 | 121178257 | 121169757 | 121169657 | 133901430 | 134049930 | 133902430 | 133901530 |
| Reference length | 119668634 | 119668634 | 119668634 | 119668634 | 119668634 | 119668634 | 119668634 | 119668634 |
| GC (%) | 36.1 | 36.1 | 36.1 | 36.1 | 36.35 | 36.35 | 36.35 | 36.35 |
| Reference GC (%) | 36.06 | 36.06 | 36.06 | 36.06 | 36.06 | 36.06 | 36.06 | 36.06 |
| N50 | 14864979 | 19600500 | 15405308 | 24256539 | 26162003 | 3413500 | 16698585 | 32656027 |
| NG50 | 14864979 | 19600500 | 15405308 | 24256539 | 30145414 | 4379420 | 17299953 | 32656027 |
| N90 | 9471025 | 12160500 | 9471025 | 19109354 | 22263862 | 175000 | 13446829 | 22674124 |
| NG90 | 9471025 | 15336445 | 9471025 | 19109354 | 22674124 | 300000 | 16050884 | 30145414 |
| auN | 13070225.3 | 21496511.8 | 14140798.4 | 24933961.5 | 27403799.5 | 6054223.9 | 18680542.8 | 36103785.1 |
| auNG | 13234069 | 21767690.8 | 14318180.5 | 25246712.2 | 30663072 | 6781796.2 | 20902470.3 | 40397821.1 |
| L50 | 4 | 3 | 4 | 3 | 3 | 8 | 4 | 2 |
| LG50 | 4 | 3 | 4 | 3 | 2 | 7 | 3 | 2 |
| L90 | 8 | 6 | 8 | 5 | 5 | 139 | 7 | 4 |
| LG90 | 8 | 5 | 8 | 5 | 4 | 82 | 6 | 3 |
| Misassemblies | 389 | 352 | 397 | 387 | 1665 | 1244 | 1689 | 1660 |
| Misassembled contigs | 14 | 20 | 13 | 5 | 5 | 25 | 8 | 4 |

Continued on next page

**Table S1 continued from previous page**

| Assembly | Flye | Flye_3d-dna | Flye_salsa | Flye_yahs | Hifiasm | Hifiasm_3d-dna | Hifiasm_salsa | Hifiasm_yahs |
|---|---|---|---|---|---|---|---|---|
| Misassembled contigs length | 121124559 | 120786675 | 121125559 | 121004459 | 133901430 | 74321589 | 133902430 | 133901530 |
| Local misassemblies | 116 | 112 | 109 | 101 | 619 | 374 | 613 | 615 |
| Scaffold gap ext. mis. | 0 | 0 | 0 | 0 | 0 | 2 | 0 | 0 |
| Scaffold gap loc. mis. | 0 | 3 | 1 | 0 | 0 | 0 | 0 | 0 |
| Unaligned mis. contigs | 1 | 2 | 1 | 2 | 0 | 29 | 0 | 0 |
| Unaligned contigs | 0 + 15 part | 0 + 14 part | 0 + 14 part | 0 + 7 part | 0 + 5 part | 8 + 60 part | 0 + 8 part | 0 + 4 part |
| Unaligned length | 1185893 | 1230774 | 1142845 | 1166251 | 11515295 | 11681283 | 11510549 | 11532188 |
| Genome fraction (%) | 99.097 | 99.068 | 99.105 | 99.109 | 99.302 | 99.304 | 99.302 | 99.287 |
| Duplication ratio | 1.011 | 1.011 | 1.011 | 1.011 | 1.027 | 1.027 | 1.027 | 1.027 |
| N's per 100 kbp | 0 | 7.84 | 0.83 | 0.74 | 0 | 110.78 | 0.75 | 0.07 |
| Mismatches per 100 kbp | 47.74 | 48.16 | 48.38 | 48.23 | 69.55 | 70.56 | 69.75 | 69.18 |
| Indels per 100 kbp | 7.93 | 8.09 | 8.03 | 7.98 | 7.97 | 7.92 | 7.97 | 7.91 |
| Genomic features | 707323 + 132 part | 707001 + 164 part | 707323 + 132 part | 707326 + 129 part | 708153 + 145 part | 706427 + 1757 part | 708159 + 145 part | 708159 + 145 part |
| Largest alignment | 9640198 | 9639526 | 9640198 | 9639526 | 9639574 | 2451962 | 9640246 | 9639574 |
| Total aligned length | 119695370 | 119690055 | 119740121 | 119721272 | 121877392 | 121849805 | 121903928 | 121892201 |
| NA50 | 3956242 | 3982976 | 3956242 | 3956242 | 3601484 | 425000 | 3601484 | 3601484 |
| NGA50 | 3956242 | 3982976 | 3956242 | 3956242 | 3956256 | 500000 | 3956256 | 3956256 |
| NA90 | 841493 | 841493 | 841493 | 841493 | 4404 | 4199 | 4404 | 4404 |
| NGA90 | 841965 | 882551 | 841965 | 841965 | 842402 | 81261 | 842402 | 842402 |
| auNA | 4310743.1 | 4325041.6 | 4309990.1 | 4312285.1 | 3899311.8 | 604328.3 | 3899704.3 | 3899858.8 |
| auNGA | 4364781.1 | 4379602.1 | 4364054.6 | 4366374.8 | 4363076.7 | 676954.1 | 4363548.5 | 4363692 |
| LA50 | 10 | 10 | 10 | 10 | 12 | 83 | 12 | 12 |
| LGA50 | 10 | 10 | 10 | 10 | 10 | 67 | 10 | 10 |
| LA90 | 37 | 36 | 37 | 37 | 317 | 870 | 316 | 313 |
| LGA90 | 36 | 35 | 36 | 36 | 36 | 284 | 36 | 36 |

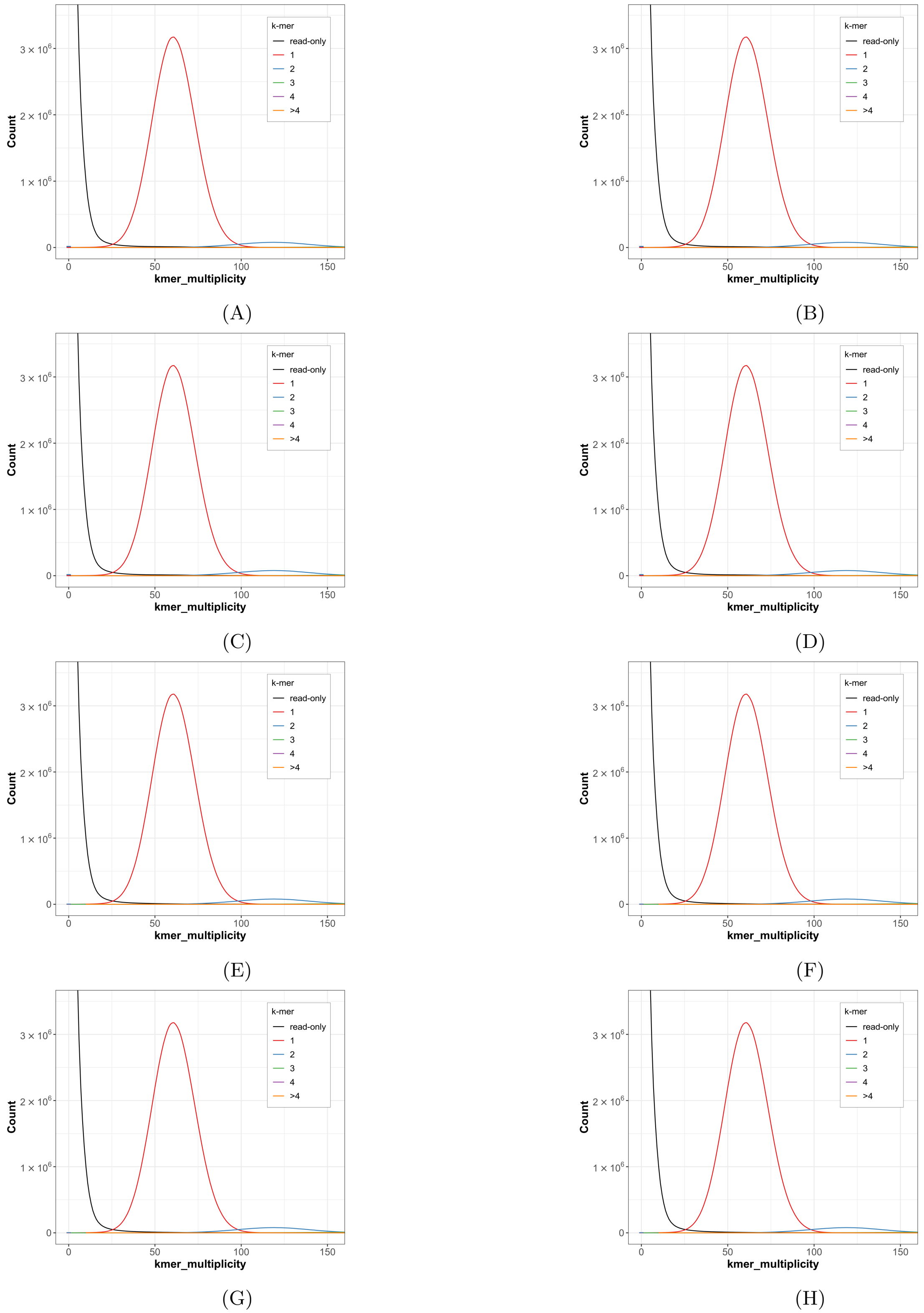


Figure S1: Copy number spectrum plots obtained with Merqury.
(A) Flye contig-level assembly, (B) Flye scaffolded with 3d-dna, (C) Flye scaffolded with SALSA2, (D) Flye scaffolded with YaHS, (E) Hifiasm contig-level assembly, (F) Hifiasm scaffolded with 3d-dna, (G) Hifiasm scaffolded with SALSA2, (H) Hifiasm scaffolded with YaHS.

Table S2: Liftoff_combine.py results for contigs or scaffolds containing genes mapping to chromosome 1 of the reference genome TAIR10.1.

| Assembly | Contigs or scaffolds corresponding to chromosome 1 | Gene count | Genes showing divergent gene distance | Reverse complement | Genes (%) in ascending order | Genes (%) in descending order | Total number of genes |
|---|---|---|---|---|---|---|---|
| Flye | contig_21_1 | 4356 | 24 | no | 99.500000 | 0.200000 | 8740.000000 |
| | contig_5_1 | 4384 | 31 | | | | |
| Flye_3d-dna | HiC_scaffold_15 | 1 | 1 | yes | 6.700000 | 93.100000 | 8740.000000 |
| | HiC_scaffold_16 | 1 | 1 | | | | |
| | HiC_scaffold_2 | 2 | 2 | | | | |
| | HiC_scaffold_5 | 8735 | 8393 | | | | |
| | HiC_scaffold_9 | 1 | 1 | | | | |
| Flye_salsa | scaffold_2 | 1 | 1 | no | 99.500000 | 0.200000 | 8740.000000 |
| | scaffold_3 | 4355 | 23 | | | | |
| | scaffold_5 | 4384 | 31 | | | | |
| Flye_yahs | scaffold_1 | 8740 | 54 | no | 99.500000 | 0.200000 | 8740.000000 |
| Hifiasm | ptg000007l_1 | 8738 | 56 | no | 99.500000 | 0.200000 | 8738.000000 |
| Hifiasm_3d-dna | HiC_scaffold_14 | 6 | 1 | undetermined | 54.300000 | 45.400000 | 8734.000000 |
| | HiC_scaffold_15 | 9 | 1 | | | | |
| | HiC_scaffold_20 | 6 | 1 | | | | |
| | HiC_scaffold_22 | 914 | 2 | | | | |
| | HiC_scaffold_23 | 88 | 1 | | | | |
| | HiC_scaffold_24 | 16 | 1 | | | | |
| | HiC_scaffold_25 | 289 | 1 | | | | |
| | HiC_scaffold_26 | 176 | 2 | | | | |
| | HiC_scaffold_27 | 81 | 1 | | | | |
| | HiC_scaffold_28 | 55 | 1 | | | | |
| | HiC_scaffold_29 | 78 | 1 | | | | |
| | HiC_scaffold_3 | 4785 | 4083 | | | | |
| | HiC_scaffold_30 | 3 | 1 | | | | |
| | HiC_scaffold_31 | 97 | 2 | | | | |
| | HiC_scaffold_32 | 166 | 2 | | | | |
| | HiC_scaffold_33 | 34 | 1 | | | | |
| | HiC_scaffold_34 | 18 | 1 | | | | |
| | HiC_scaffold_35 | 86 | 2 | | | | |
| | HiC_scaffold_36 | 13 | 7 | | | | |
| | HiC_scaffold_37 | 37 | 1 | | | | |
| | HiC_scaffold_38 | 17 | 1 | | | | |
| | HiC_scaffold_39 | 65 | 2 | | | | |
| | HiC_scaffold_40 | 53 | 1 | | | | |

Continued on next page

**Table S2 continued from previous page**

| Assembly | Contigs or scaffolds corresponding to chromosome 1 | Gene count | Genes showing divergent gene distance | Reverse complement | Genes (%) in ascending order | Genes (%) in descending order | Total number of genes |
|---|---|---|---|---|---|---|---|
| | HiC_scaffold_41 | 31 | 1 | | | | |
| | HiC_scaffold_42 | 27 | 1 | | | | |
| | HiC_scaffold_43 | 86 | 1 | | | | |
| | HiC_scaffold_44 | 86 | 1 | | | | |
| | HiC_scaffold_45 | 15 | 1 | | | | |
| | HiC_scaffold_46 | 22 | 1 | | | | |
| | HiC_scaffold_47 | 19 | 1 | | | | |
| | HiC_scaffold_48 | 34 | 1 | | | | |
| | HiC_scaffold_49 | 21 | 1 | | | | |
| | HiC_scaffold_50 | 7 | 1 | | | | |
| | HiC_scaffold_51 | 9 | 1 | | | | |
| | HiC_scaffold_68 | 40 | 1 | | | | |
| | HiC_scaffold_69 | 38 | 1 | | | | |
| | HiC_scaffold_70 | 68 | 1 | | | | |
| | HiC_scaffold_71 | 27 | 1 | | | | |
| | HiC_scaffold_72 | 43 | 1 | | | | |
| | HiC_scaffold_73 | 39 | 1 | | | | |
| | HiC_scaffold_74 | 14 | 1 | | | | |
| | HiC_scaffold_75 | 12 | 1 | | | | |
| | HiC_scaffold_76 | 8 | 1 | | | | |
| | HiC_scaffold_77 | 86 | 1 | | | | |
| | HiC_scaffold_78 | 56 | 1 | | | | |
| | HiC_scaffold_79 | 3 | 1 | | | | |
| | HiC_scaffold_80 | 7 | 1 | | | | |
| | HiC_scaffold_81 | 16 | 1 | | | | |
| | HiC_scaffold_82 | 40 | 1 | | | | |
| | HiC_scaffold_83 | 33 | 1 | | | | |
| | HiC_scaffold_84 | 9 | 1 | | | | |
| | HiC_scaffold_85 | 59 | 1 | | | | |
| | HiC_scaffold_86 | 12 | 1 | | | | |
| | HiC_scaffold_87 | 8 | 1 | | | | |
| | HiC_scaffold_88 | 13 | 1 | | | | |
| | HiC_scaffold_89 | 10 | 1 | | | | |
| | HiC_scaffold_90 | 101 | 2 | | | | |
| | HiC_scaffold_91 | 36 | 1 | | | | |
| | HiC_scaffold_92 | 87 | 1 | | | | |
| | HiC_scaffold_93 | 44 | 1 | | | | |

Continued on next page

**Table S2 continued from previous page**

| Assembly | Contigs or scaffolds corresponding to chromosome 1 | Gene count | Genes showing divergent gene distance | Reverse complement | Genes (%) in ascending order | Genes (%) in descending order | Total number of genes |
|---|---|---|---|---|---|---|---|
| | HiC_scaffold_94 | 57 | 1 | | | | |
| | HiC_scaffold_95 | 130 | 1 | | | | |
| | HiC_scaffold_96 | 127 | 1 | | | | |
| | HiC_scaffold_97 | 62 | 1 | | | | |
| Hifiasm_salsa | scaffold_5 | 4356 | 24 | no | 99.500000 | 0.200000 | 8738.000000 |
| | scaffold_6 | 4382 | 32 | | | | |
| Hifiasm_yahs | scaffold_2 | 8738 | 55 | no | 99.500000 | 0.200000 | 8738.000000 |

Table S3: Liftoff_combine.py results for contigs or scaffolds containing genes mapping to chromosome 2 of the reference genome TAIR10.1.

| Assembly | Contigs or scaffolds corresponding to chromosome 2 | Gene count | Genes showing divergent gene distance | Reverse complement | Genes (%) in ascending order | Genes (%) in descending order | Total number of genes |
|---|---|---|---|---|---|---|---|
| Flye | contig_16_1 | 1 | 1 | yes | 6.100000 | 93.700000 | 5190.000000 |
| | contig_1_1 | 724 | 696 | | | | |
| | contig_21_1 | 1 | 1 | | | | |
| | contig_27_1 | 1 | 1 | | | | |
| | contig_31_1 | 2 | 2 | | | | |
| | contig_4_1 | 4458 | 4294 | | | | |
| | contig_5_1 | 1 | 1 | | | | |
| | contig_66_1 | 1 | 1 | | | | |
| | contig_6_1 | 1 | 1 | | | | |
| Flye_3d-dna | HiC_scaffold_1 | 2 | 2 | undetermined | 86.600000 | 13.100000 | 5192.000000 |
| | HiC_scaffold_2 | 3 | 3 | | | | |
| | HiC_scaffold_22 | 22 | 19 | | | | |
| | HiC_scaffold_23 | 4 | 4 | | | | |
| | HiC_scaffold_24 | 3 | 2 | | | | |
| | HiC_scaffold_3 | 1 | 1 | | | | |
| | HiC_scaffold_4 | 5155 | 680 | | | | |
| | HiC_scaffold_5 | 2 | 2 | | | | |
| Flye_salsa | scaffold_13 | 2 | 2 | undetermined | 86.600000 | 13.100000 | 5191.000000 |
| | scaffold_2 | 4459 | 9 | | | | |
| | scaffold_3 | 1 | 1 | | | | |
| | scaffold_4 | 1 | 1 | | | | |

Continued on next page

**Table S3 continued from previous page**

| Assembly | Contigs or scaffolds corresponding to chromosome 2 | Gene count | Genes showing divergent gene distance | Reverse complement | Genes (%) in ascending order | Genes (%) in descending order | Total number of genes |
|---|---|---|---|---|---|---|---|
| | scaffold_5 | 1 | 1 | | | | |
| | scaffold_6 | 1 | 1 | | | | |
| | scaffold_7 | 1 | 1 | | | | |
| | scaffold_8 | 1 | 1 | | | | |
| | scaffold_9 | 724 | 696 | | | | |
| Flye_yahs | scaffold_1 | 2 | 2 | yes | 6.000000 | 93.800000 | 5192.000000 |
| | scaffold_2 | 2 | 2 | | | | |
| | scaffold_3 | 4 | 4 | | | | |
| | scaffold_4 | 5184 | 4990 | | | | |
| Hifiasm | ptg000001l_1 | 5260 | 5064 | yes | 5.900000 | 93.900000 | 5260.000000 |
| Hifiasm_3d-dna | HiC_scaffold_1 | 809 | 29 | undetermined | 44.600000 | 55.100000 | 5259.000000 |
| | HiC_scaffold_2 | 1664 | 325 | | | | |
| | HiC_scaffold_240 | 175 | 166 | | | | |
| | HiC_scaffold_241 | 204 | 193 | | | | |
| | HiC_scaffold_242 | 231 | 225 | | | | |
| | HiC_scaffold_243 | 447 | 429 | | | | |
| | HiC_scaffold_244 | 20 | 20 | | | | |
| | HiC_scaffold_245 | 55 | 52 | | | | |
| | HiC_scaffold_246 | 84 | 83 | | | | |
| | HiC_scaffold_247 | 85 | 82 | | | | |
| | HiC_scaffold_248 | 94 | 90 | | | | |
| | HiC_scaffold_249 | 25 | 24 | | | | |
| | HiC_scaffold_250 | 37 | 35 | | | | |
| | HiC_scaffold_251 | 156 | 142 | | | | |
| | HiC_scaffold_252 | 96 | 89 | | | | |
| | HiC_scaffold_253 | 192 | 185 | | | | |
| | HiC_scaffold_254 | 71 | 68 | | | | |
| | HiC_scaffold_255 | 27 | 26 | | | | |
| | HiC_scaffold_256 | 46 | 45 | | | | |
| | HiC_scaffold_257 | 69 | 67 | | | | |
| | HiC_scaffold_258 | 22 | 22 | | | | |
| | HiC_scaffold_259 | 6 | 6 | | | | |
| | HiC_scaffold_260 | 83 | 81 | | | | |
| | HiC_scaffold_261 | 6 | 6 | | | | |
| | HiC_scaffold_262 | 276 | 262 | | | | |
| | HiC_scaffold_263 | 36 | 36 | | | | |

Continued on next page

**Table S3 continued from previous page**

| Assembly | Contigs or scaffolds corresponding to chromosome 2 | Gene count | Genes showing divergent gene distance | Reverse complement | Genes (%) in ascending order | Genes (%) in descending order | Total number of genes |
|---|---|---|---|---|---|---|---|
| | HiC_scaffold_264 | 125 | 120 | | | | |
| | HiC_scaffold_265 | 10 | 9 | | | | |
| | HiC_scaffold_266 | 7 | 7 | | | | |
| | HiC_scaffold_268 | 3 | 3 | | | | |
| | HiC_scaffold_269 | 3 | 3 | | | | |
| | HiC_scaffold_279 | 63 | 60 | | | | |
| | HiC_scaffold_280 | 24 | 22 | | | | |
| | HiC_scaffold_282 | 1 | 1 | | | | |
| | HiC_scaffold_283 | 4 | 4 | | | | |
| | HiC_scaffold_342 | 3 | 3 | | | | |
| Hifiasm_salsa | scaffold_3 | 4461 | 11 | undetermined | 85.500000 | 14.300000 | 5261.000000 |
| | scaffold_8 | 800 | 768 | | | | |
| Hifiasm_yahs | scaffold_4 | 5260 | 5064 | yes | 5.900000 | 93.900000 | 5261.000000 |

Table S4: Liftoff_combine.py results for contigs or scaffolds containing genes mapping to chromosome 3 of the reference genome TAIR10.1.

| Assembly | Contigs or scaffolds corresponding to chromosome 3 | Gene count | Genes showing divergent gene distance | Reverse complement | Genes (%) in ascending order | Genes (%) in descending order | Total number of genes |
|---|---|---|---|---|---|---|---|
| Flye | contig_11_1 | 10 | 5 | yes | 5.900000 | 93.900000 | 6543.000000 |
| | contig_27_1 | 2650 | 2565 | | | | |
| | contig_31_1 | 8 | 8 | | | | |
| | contig_6_1 | 3875 | 3748 | | | | |
| Flye_3d-dna | HiC_scaffold_1 | 6533 | 14 | no | 99.600000 | 0.100000 | 6543.000000 |
| | HiC_scaffold_26 | 3 | 2 | | | | |
| | HiC_scaffold_6 | 7 | 5 | | | | |
| Flye_salsa | scaffold_1 | 10 | 7 | yes | 5.900000 | 93.900000 | 6543.000000 |
| | scaffold_12 | 8 | 8 | | | | |
| | scaffold_6 | 3875 | 3748 | | | | |
| | scaffold_8 | 2650 | 2565 | | | | |
| Flye_yahs | scaffold_3 | 6533 | 14 | no | 99.700000 | 0.100000 | 6543.000000 |
| | scaffold_5 | 10 | 5 | | | | |
| Hifiasm | ptg000001l_1 | 7 | 7 | no | 99.600000 | 0.100000 | 6542.000000 |
| | ptg000002l_1 | 2 | 2 | | | | |

Continued on next page

**Table S4 continued from previous page**

| Assembly | Contigs or scaffolds corresponding to chromosome 3 | Gene count | Genes showing divergent gene distance | Reverse complement | Genes (%) in ascending order | Genes (%) in descending order | Total number of genes |
|---|---|---|---|---|---|---|---|
| | ptg000003l_1 | 6533 | 14 | | | | |
| Hifiasm_3d-dna | HiC_scaffold_17 | 5 | 1 | undetermined | 76.081970 | 23.673345 | 6540.000000 |
| | HiC_scaffold_173 | 251 | 1 | | | | |
| | HiC_scaffold_174 | 177 | 1 | | | | |
| | HiC_scaffold_175 | 94 | 1 | | | | |
| | HiC_scaffold_176 | 177 | 1 | | | | |
| | HiC_scaffold_177 | 118 | 1 | | | | |
| | HiC_scaffold_178 | 150 | 1 | | | | |
| | HiC_scaffold_179 | 103 | 1 | | | | |
| | HiC_scaffold_18 | 7 | 1 | | | | |
| | HiC_scaffold_180 | 180 | 1 | | | | |
| | HiC_scaffold_181 | 145 | 3 | | | | |
| | HiC_scaffold_182 | 200 | 1 | | | | |
| | HiC_scaffold_183 | 32 | 1 | | | | |
| | HiC_scaffold_184 | 149 | 1 | | | | |
| | HiC_scaffold_185 | 81 | 1 | | | | |
| | HiC_scaffold_186 | 17 | 1 | | | | |
| | HiC_scaffold_187 | 176 | 1 | | | | |
| | HiC_scaffold_188 | 181 | 1 | | | | |
| | HiC_scaffold_189 | 7 | 1 | | | | |
| | HiC_scaffold_19 | 3 | 1 | | | | |
| | HiC_scaffold_190 | 19 | 1 | | | | |
| | HiC_scaffold_191 | 6 | 1 | | | | |
| | HiC_scaffold_192 | 23 | 1 | | | | |
| | HiC_scaffold_193 | 30 | 1 | | | | |
| | HiC_scaffold_194 | 61 | 1 | | | | |
| | HiC_scaffold_195 | 6 | 1 | | | | |
| | HiC_scaffold_196 | 48 | 1 | | | | |
| | HiC_scaffold_197 | 26 | 1 | | | | |
| | HiC_scaffold_198 | 31 | 1 | | | | |
| | HiC_scaffold_199 | 40 | 1 | | | | |
| | HiC_scaffold_200 | 66 | 1 | | | | |
| | HiC_scaffold_201 | 3 | 1 | | | | |
| | HiC_scaffold_202 | 2 | 1 | | | | |
| | HiC_scaffold_213 | 1 | 1 | | | | |
| | HiC_scaffold_214 | 1 | 1 | | | | |
| | HiC_scaffold_215 | 24 | 2 | | | | |

Continued on next page

**Table S4 continued from previous page**

| Assembly | Contigs or scaffolds corresponding to chromosome 3 | Gene count | Genes showing divergent gene distance | Reverse complement | Genes (%) in ascending order | Genes (%) in descending order | Total number of genes |
|---|---|---|---|---|---|---|---|
| | HiC_scaffold_216 | 2 | 1 | | | | |
| | HiC_scaffold_217 | 14 | 1 | | | | |
| | HiC_scaffold_218 | 46 | 1 | | | | |
| | HiC_scaffold_219 | 8 | 1 | | | | |
| | HiC_scaffold_220 | 7 | 1 | | | | |
| | HiC_scaffold_221 | 91 | 1 | | | | |
| | HiC_scaffold_222 | 77 | 2 | | | | |
| | HiC_scaffold_223 | 26 | 1 | | | | |
| | HiC_scaffold_224 | 69 | 1 | | | | |
| | HiC_scaffold_225 | 159 | 1 | | | | |
| | HiC_scaffold_226 | 138 | 1 | | | | |
| | HiC_scaffold_227 | 108 | 1 | | | | |
| | HiC_scaffold_228 | 215 | 1 | | | | |
| | HiC_scaffold_229 | 63 | 1 | | | | |
| | HiC_scaffold_230 | 63 | 1 | | | | |
| | HiC_scaffold_231 | 224 | 1 | | | | |
| | HiC_scaffold_232 | 82 | 1 | | | | |
| | HiC_scaffold_233 | 277 | 1 | | | | |
| | HiC_scaffold_234 | 7 | 1 | | | | |
| | HiC_scaffold_235 | 22 | 1 | | | | |
| | HiC_scaffold_236 | 43 | 1 | | | | |
| | HiC_scaffold_237 | 66 | 1 | | | | |
| | HiC_scaffold_238 | 59 | 1 | | | | |
| | HiC_scaffold_239 | 125 | 1 | | | | |
| | HiC_scaffold_284 | 7 | 6 | | | | |
| | HiC_scaffold_342 | 1 | 1 | | | | |
| | HiC_scaffold_6 | 1900 | 1561 | | | | |
| | HiC_scaffold_8 | 1 | 1 | | | | |
| Hifiasm_salsa | scaffold_1 | 6533 | 6321 | yes | 5.900000 | 93.900000 | 6542.000000 |
| | scaffold_2 | 2 | 2 | | | | |
| | scaffold_8 | 7 | 6 | | | | |
| Hifiasm_yahs | scaffold_1 | 6535 | 16 | no | 99.600000 | 0.100000 | 6542.000000 |
| | scaffold_4 | 7 | 6 | | | | |

12

Table S5: Liftoff_combine.py results for contigs or scaffolds containing genes mapping to chromosome 4 of the reference genome TAIR10.1.

| Assembly | Contigs or scaffolds corresponding to chromosome 4 | Gene count | Genes showing divergent gene distance | Reverse complement | Genes (%) in ascending order | Genes (%) in descending order | Total number of genes |
|---|---|---|---|---|---|---|---|
| Flye | contig_11_1 | 737 | 4 | no | 98.800000 | 1.000000 | 4998.000000 |
| | contig_14_1 | 17 | 3 | | | | |
| | contig_18_1 | 4200 | 21 | | | | |
| | contig_67_1 | 44 | 42 | | | | |
| Flye_3d-dna | HiC_scaffold_20 | 1 | 1 | yes | 5.500000 | 94.200000 | 4998.000000 |
| | HiC_scaffold_6 | 4997 | 4814 | | | | |
| Flye_salsa | scaffold_1 | 4954 | 746 | undetermined | 84.900000 | 14.900000 | 4998.000000 |
| | scaffold_10 | 44 | 42 | | | | |
| Flye_yahs | scaffold_5 | 4998 | 31 | no | 99.600000 | 0.100000 | 4998.000000 |
| Hifiasm | ptg000002l_1 | 5000 | 4818 | yes | 5.500000 | 94.200000 | 5000.000000 |
| Hifiasm_3d-dna | HiC_scaffold_285 | 293 | 278 | yes | 9.700000 | 90.000000 | 4999.000000 |
| | HiC_scaffold_286 | 9 | 9 | | | | |
| | HiC_scaffold_287 | 75 | 72 | | | | |
| | HiC_scaffold_288 | 7 | 7 | | | | |
| | HiC_scaffold_289 | 223 | 213 | | | | |
| | HiC_scaffold_290 | 103 | 100 | | | | |
| | HiC_scaffold_291 | 155 | 148 | | | | |
| | HiC_scaffold_292 | 120 | 116 | | | | |
| | HiC_scaffold_293 | 241 | 238 | | | | |
| | HiC_scaffold_294 | 98 | 95 | | | | |
| | HiC_scaffold_295 | 213 | 208 | | | | |
| | HiC_scaffold_296 | 283 | 275 | | | | |
| | HiC_scaffold_297 | 17 | 17 | | | | |
| | HiC_scaffold_298 | 135 | 124 | | | | |
| | HiC_scaffold_299 | 82 | 79 | | | | |
| | HiC_scaffold_300 | 34 | 34 | | | | |
| | HiC_scaffold_301 | 21 | 19 | | | | |
| | HiC_scaffold_302 | 30 | 29 | | | | |
| | HiC_scaffold_303 | 7 | 7 | | | | |
| | HiC_scaffold_304 | 13 | 13 | | | | |
| | HiC_scaffold_305 | 21 | 21 | | | | |
| | HiC_scaffold_306 | 68 | 67 | | | | |
| | HiC_scaffold_307 | 16 | 15 | | | | |
| | HiC_scaffold_308 | 55 | 53 | | | | |
| | HiC_scaffold_309 | 36 | 35 | | | | |

Continued on next page

**Table S5 continued from previous page**

| Assembly | Contigs or scaffolds corresponding to chromosome 4 | Gene count | Genes showing divergent gene distance | Reverse complement | Genes (%) in ascending order | Genes (%) in descending order | Total number of genes |
|---|---|---|---|---|---|---|---|
| | HiC_scaffold_310 | 32 | 31 | | | | |
| | HiC_scaffold_311 | 146 | 140 | | | | |
| | HiC_scaffold_312 | 28 | 27 | | | | |
| | HiC_scaffold_313 | 103 | 102 | | | | |
| | HiC_scaffold_314 | 95 | 95 | | | | |
| | HiC_scaffold_315 | 42 | 41 | | | | |
| | HiC_scaffold_326 | 3 | 3 | | | | |
| | HiC_scaffold_328 | 12 | 12 | | | | |
| | HiC_scaffold_329 | 15 | 15 | | | | |
| | HiC_scaffold_330 | 14 | 13 | | | | |
| | HiC_scaffold_331 | 7 | 7 | | | | |
| | HiC_scaffold_332 | 10 | 10 | | | | |
| | HiC_scaffold_333 | 7 | 6 | | | | |
| | HiC_scaffold_334 | 6 | 5 | | | | |
| | HiC_scaffold_336 | 54 | 54 | | | | |
| | HiC_scaffold_337 | 16 | 16 | | | | |
| | HiC_scaffold_338 | 8 | 7 | | | | |
| | HiC_scaffold_339 | 31 | 31 | | | | |
| | HiC_scaffold_340 | 33 | 31 | | | | |
| | HiC_scaffold_341 | 98 | 95 | | | | |
| | HiC_scaffold_7 | 1252 | 1068 | | | | |
| | HiC_scaffold_8 | 632 | 579 | | | | |
| Hifiams_salsa | scaffold_2 | 5000 | 750 | undetermined | 85.600000 | 14.100000 | 5000.000000 |
| Hifiasm_yahs | scaffold_1 | 5000 | 33 | no | 99.600000 | 0.200000 | 5000.000000 |

Table S6: Liftoff_combine.py results for contigs or scaffolds containing genes mapping to chromosome 5 of the reference genome TAIR10.1.

| Assembly | Contigs or scaffolds corresponding to chromosome 5 | Gene count | Genes showing divergent gene distance | Reverse complement | Genes (%) in ascending order | Genes (%) in descending order | Total number of genes |
|---|---|---|---|---|---|---|---|
| Flye | contig_13_1 | 11 | 1 | undetermined | 46.7 | 53.1 | 7449 |
| | contig_16_1 | 4148 | 4034 | | | | |
| | contig_66_1 | 3290 | 7 | | | | |
| Flye_3d-dna | HiC_scaffold_2 | 3301 | 3198 | yes | 5 | 94.9 | 7449 |

Continued on next page

**Table S6 continued from previous page**

| Assembly | Contigs or scaffolds corresponding to chromosome 5 | Gene count | Genes showing divergent gene distance | Reverse complement | Genes (%) in ascending order | Genes (%) in descending order | Total number of genes |
|---|---|---|---|---|---|---|---|
| | HiC_scaffold_21 | 1 | 1 | | | | |
| | HiC_scaffold_3 | 4147 | 4033 | | | | |
| Flye_salsa | scaffold_11 | 11 | 1 | undetermined | 46.7 | 53.1 | 7449 |
| | scaffold_4 | 4148 | 4034 | | | | |
| | scaffold_7 | 3290 | 7 | | | | |
| Flye_yahs | scaffold_2 | 7449 | 7222 | yes | 5.1 | 94.7 | 7449 |
| Hifiasm | ptg000006l_1 | 7447 | 37 | no | 99.7 | 0.1 | 7447 |
| Hifiasm_3d-dna | HiC_scaffold_1 | 1 | 1 | undetermined | 69 | 30.8 | 7443 |
| | HiC_scaffold_100 | 545 | 2 | | | | |
| | HiC_scaffold_101 | 80 | 1 | | | | |
| | HiC_scaffold_102 | 23 | 1 | | | | |
| | HiC_scaffold_103 | 39 | 1 | | | | |
| | HiC_scaffold_104 | 109 | 1 | | | | |
| | HiC_scaffold_105 | 105 | 2 | | | | |
| | HiC_scaffold_106 | 110 | 1 | | | | |
| | HiC_scaffold_107 | 43 | 1 | | | | |
| | HiC_scaffold_108 | 183 | 1 | | | | |
| | HiC_scaffold_109 | 146 | 1 | | | | |
| | HiC_scaffold_110 | 62 | 1 | | | | |
| | HiC_scaffold_111 | 39 | 1 | | | | |
| | HiC_scaffold_112 | 71 | 2 | | | | |
| | HiC_scaffold_113 | 140 | 1 | | | | |
| | HiC_scaffold_114 | 12 | 1 | | | | |
| | HiC_scaffold_115 | 68 | 1 | | | | |
| | HiC_scaffold_116 | 172 | 1 | | | | |
| | HiC_scaffold_117 | 13 | 1 | | | | |
| | HiC_scaffold_118 | 14 | 1 | | | | |
| | HiC_scaffold_119 | 16 | 1 | | | | |
| | HiC_scaffold_120 | 73 | 1 | | | | |
| | HiC_scaffold_121 | 14 | 1 | | | | |
| | HiC_scaffold_122 | 20 | 1 | | | | |
| | HiC_scaffold_123 | 20 | 1 | | | | |
| | HiC_scaffold_124 | 20 | 1 | | | | |
| | HiC_scaffold_125 | 17 | 1 | | | | |
| | HiC_scaffold_126 | 10 | 1 | | | | |
| | HiC_scaffold_134 | 7 | 1 | | | | |

Continued on next page

**Table S6 continued from previous page**

| Assembly | Contigs or scaffolds corresponding to chromosome 5 | Gene count | Genes showing divergent gene distance | Reverse complement | Genes (%) in ascending order | Genes (%) in descending order | Total number of genes |
|---|---|---|---|---|---|---|---|
| | HiC_scaffold_135 | 1 | 1 | | | | |
| | HiC_scaffold_136 | 3 | 1 | | | | |
| | HiC_scaffold_137 | 2 | 1 | | | | |
| | HiC_scaffold_138 | 3 | 1 | | | | |
| | HiC_scaffold_139 | 7 | 1 | | | | |
| | HiC_scaffold_140 | 53 | 1 | | | | |
| | HiC_scaffold_141 | 1 | 1 | | | | |
| | HiC_scaffold_142 | 23 | 1 | | | | |
| | HiC_scaffold_143 | 15 | 1 | | | | |
| | HiC_scaffold_144 | 28 | 1 | | | | |
| | HiC_scaffold_145 | 70 | 1 | | | | |
| | HiC_scaffold_146 | 49 | 1 | | | | |
| | HiC_scaffold_147 | 89 | 1 | | | | |
| | HiC_scaffold_148 | 9 | 1 | | | | |
| | HiC_scaffold_149 | 25 | 1 | | | | |
| | HiC_scaffold_150 | 30 | 1 | | | | |
| | HiC_scaffold_151 | 17 | 1 | | | | |
| | HiC_scaffold_152 | 22 | 1 | | | | |
| | HiC_scaffold_153 | 35 | 1 | | | | |
| | HiC_scaffold_154 | 65 | 1 | | | | |
| | HiC_scaffold_155 | 32 | 2 | | | | |
| | HiC_scaffold_156 | 82 | 1 | | | | |
| | HiC_scaffold_157 | 97 | 1 | | | | |
| | HiC_scaffold_158 | 63 | 1 | | | | |
| | HiC_scaffold_159 | 110 | 1 | | | | |
| | HiC_scaffold_160 | 52 | 1 | | | | |
| | HiC_scaffold_161 | 11 | 1 | | | | |
| | HiC_scaffold_162 | 67 | 1 | | | | |
| | HiC_scaffold_163 | 158 | 1 | | | | |
| | HiC_scaffold_164 | 69 | 1 | | | | |
| | HiC_scaffold_165 | 233 | 1 | | | | |
| | HiC_scaffold_166 | 96 | 1 | | | | |
| | HiC_scaffold_167 | 33 | 1 | | | | |
| | HiC_scaffold_168 | 140 | 1 | | | | |
| | HiC_scaffold_169 | 26 | 1 | | | | |
| | HiC_scaffold_170 | 204 | 1 | | | | |
| | HiC_scaffold_171 | 124 | 2 | | | | |

Continued on next page

**Table S6 continued from previous page**

| Assembly | Contigs or scaffolds corresponding to chromosome 5 | Gene count | Genes showing divergent gene distance | Reverse complement | Genes (%) in ascending order | Genes (%) in descending order | Total number of genes |
|---|---|---|---|---|---|---|---|
| | HiC_scaffold_172 | 268 | 21 | | | | |
| | HiC_scaffold_2 | 22 | 1404 | | | | |
| | HiC_scaffold_4 | 1682 | 36 | | | | |
| | HiC_scaffold_5 | 48 | 871 | | | | |
| | HiC_scaffold_6 | 1000 | 1 | | | | |
| | HiC_scaffold_98 | 33 | 1 | | | | |
| | HiC_scaffold_99 | 74 | | | | | |
| Hifiasm_salsa | scaffold_4 | 4146 | 27 | no | 99.7 | 0.1 | 7447 |
| | scaffold_7 | 3301 | 9 | | | | |
| Hifiasm_yahs | scaffold_3 | 7447 | 37 | no | 99.7 | 0.1 | 7447 |
